# *Ex vivo* astrocyte-to-oligodendrocyte conversion in human adult cortical tissue using transcription factor overexpression

**DOI:** 10.64898/2026.03.14.711766

**Authors:** Ayesha Prajapati, Laura R. Rodríguez, Raquel Martinez-Curiel, Kenia Esparza Ocampo, Wendy P. Gastélum Espinoza, Henrik Ahlenius, Johan Bengzon, Sara Palma-Tortosa

## Abstract

Multiple sclerosis (MS) is an autoimmune and neurological disorder characterized by myelin disruption and neuronal degeneration. Currently approved therapies focus on symptom relief but do not promote central nervous system (CNS) repair. In contrast, astrocytes proliferate and repopulate MS-related lesions. Moreover, in active lesions, they hinder regenerative processes such as neural progenitor migration. Here, we propose astrocytes as a potential target for myelin repair in the human diseased brain. To achieve this aim, we investigated whether glial fibrillary acidic protein (GFAP)+ astrocytes can be transdifferentiated into oligodendrocyte lineage cells through forced overexpression of transcription factors both *in vitro* and *ex vivo* organotypic cultures of human adult cortex. Our results show that overexpression of OLIG2 and SOX10 in human induced pluripotent stem cell-derived astrocytes gives rise to oligodendrocyte progenitor cells 12 days post-induction, as shown by morphological changes and O4 marker expression. Importantly, transdifferentiation of GFAP-expressing endogenous astrocytes in human adult cortical tissue give rise to mature oligodendrocytes, as shown by expression of CC1, after only 12 days of overexpression of OLIG2 and SOX10. To our knowledge, this is the first study to assess direct astrocyte-to-oligodendrocyte reprogramming in a human platform preserving the native three-dimensional architecture of the brain. Further work will be required to determine whether the reprogrammed cells can myelinate axons and to evaluate the potential of this approach for structural and functional repair in the demyelinated human CNS.

## INTRODUCTION

Multiple sclerosis (MS) currently affects nearly 3 million people worldwide. In this condition, autoimmune-mediated inflammation targets the central nervous system (CNS), causing the loss or dysfunction of oligodendrocytes, the cells responsible for producing myelin. These events alter the remyelination process, resulting in progressive axonal injury, neuronal damage and ultimately contributing to motor, sensory and cognitive deficits. Currently, available treatments for MS are only partially effective at preventing tissue damage. Furthermore, none of them promote CNS repair, which is crucial for recovery (Tintore, Vidal-Jordana, and Sastre-Garriga 2019).

Astrocytes are glial cells that are more resilient to CNS injury than neurons and oligodendrocytes. They undergo profound morphological and transcriptional changes in response to pathology (Pekny et al. 2016). In MS, astrocytes proliferate and expand within demyelinated lesions, forming a glial scar in response to oligodendrocyte loss and myelin damage (Ponath, Park, and Pitt 2018). This reactive astrogliosis serves a dual role: in the early stages, astrocytes support lesion repair by recruiting microglia to eliminate myelin debris and by releasing neuroprotective factors, thereby facilitating remyelination. However, during chronic demyelination, prolonged astrocyte reactivity becomes detrimental, as the glial scar limits progenitor cell migration and reactive astrocytes produce pro-inflammatory cytokines, reactive oxygen species, and neurotoxic mediators that exacerbate tissue damage (Healy et al. 2022). Given their prevalence and strategic position in MS lesions, astrocytes represent a promising therapeutic target for therapies aimed at promoting myelin regeneration.

Direct conversion of human astrocytes into functional neurons has been achieved (Talifu et al. 2023). However, no effective protocols for the conversion of human astrocytes into myelinating oligodendrocytes have been published to date. *In vivo* direct lineage conversion of rodent resident astrocytes into functional oligodendrocytes has been demonstrated using either a combination of microRNA (miR-302/367) and the chromatin modifier valproate (Ghasemi-Kasman et al. 2018), or transcription factor overexpression, including Sox2 (Farhangi et al. 2019) and Sox10 (Mokhtarzadeh Khanghahi et al. 2018)(Bajohr et al., 2025).

Regarding cells of human origin, Zare and collaborators have demonstrated that a human astrocytoma cell line can be reprogrammed *in vitro* into oligodendrocyte lineage cells using epigenetic modifiers (Zare, Baharvand, and Javan 2018, 2019). Additionally, a combination of six small molecules was found to be effective in producing cells from the oligodendrocyte lineage from two human astrocyte lines, acquiring oligodendrocyte progenitor cell (OPC) morphology 5 days post-induction and expressing OPC markers after 15 days and mature oligodendrocyte markers after 30 days post-induction (Sharifi-Kelishadi et al. 2024). Also, they survived and maintained their phenotype after transplantation into demyelinated animals.

As reported, previous efforts to reprogram human astrocytes towards the oligodendrocyte lineage have been limited to *in vitro* systems or xenotransplantation models. The direct conversion of human astrocytes into oligodendrocytes within a human 3D brain microenvironment has never been explored. Here, we investigated whether overexpression of the transcription factors SOX10 and OLIG2 can induce the *in situ* conversion of astrocytes into oligodendrocytes in human brain tissue.

## RESULTS

### Overexpression of SOX10 and OLIG2 in human iPS cell-derived astrocytes induces the generation of oligodendrocytes *in vitro*

Astrocytes derived from human induced pluripotent stem (iPS)-cells were generated using the protocol described by Canals and collaborators (Canals et al. 2018). The resulting cells displayed astrocytic morphology and expressed the lineage-specific markers glial fibrillary acidic protein (GFAP) and Vimentin, as shown in **Supplementary Figure 1A**,**B**. Given that OLIG2 and SOX10 are key transcription factors involved in oligodendrocyte specification (Pozniak et al. 2010; Lu et al. 2002; Zhou and Anderson 2002; Liu et al. 2007), we tested if their overexpression would be sufficient to transdifferentiate astrocytes into an oligodendroglial fate *in vitro* (**Figure 1A**). To address this, we generated two independent constructs in which the transcription factors OLIG2 and SOX10 were driven by the GFAP promoter, ensuring selective expression within the astrocytic population. To be able to track the transduced cells, constructs included the fluorescent reporters GFP and mCherry (**Figure 1B**).

**Figure 1.**
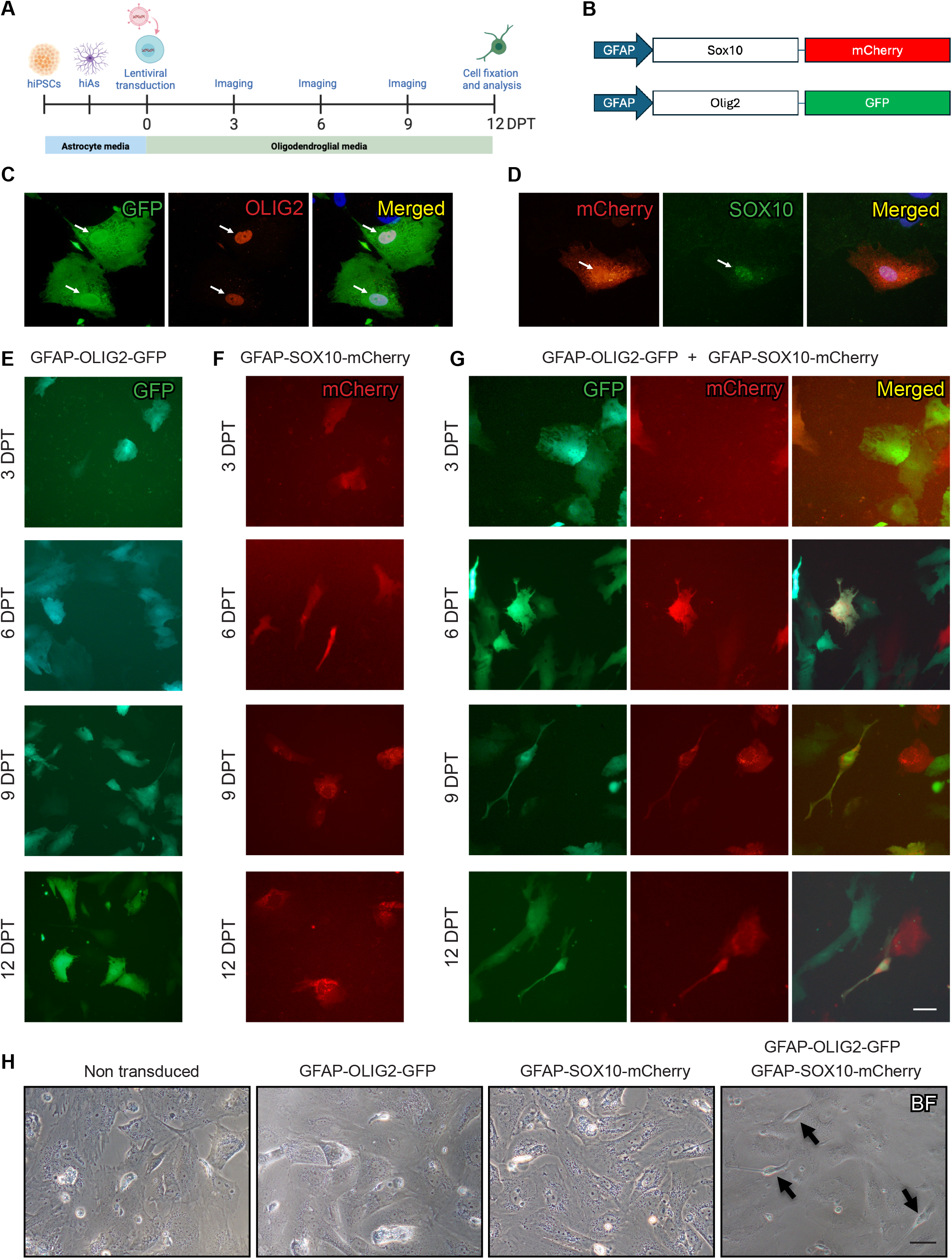
Timeline of the morphological changes induced by OLIG2 and SOX10 overexpression in human induced pluripotent stem (iPS) cell-derived astrocytes. (**A**) Experimental workflow for generating oligodendrocytes from human iPS cell-derived astrocytes. (**B**) Lentiviral vectors used for transcription factor-mediated transdifferentiation. (**C**,**D**) Representative confocal images showing colocalization of OLIG2 with the GFP reporter (**C**) and SOX10 with the mCherry reporter (**D**). (**E**-**G**). Representative fluorescence images of live cells 3, 6, 9 and 12 days post-transduction with GFAP-OLIG2-GFP (**E**), GFAP-SOX10-mCherry (**F**) or a combination of both lentiviruses (**G**). Reporter fluorescence (GFP and mCherry) is shown. (**H**) Brightfield images of the different culture conditions 12 days post-transduction. Arrows indicate cells exhibiting morphological changes. Scale bars, 50 μm.

Human iPS cell-derived astrocytes were transduced with either GFAP-OLIG2-GFP, GFAP-SOX10-mCherry or a combination of both lentiviruses. Non-transduced astrocytes were used as a control. Morphological changes were monitored in culture for up to day 12 of induction (**Figure 1**).

We first validated the lentiviral delivery *in vitro* by confirming the expression of OLIG2 in GFP-positive cells and SOX10 in mCherry-positive cells (**Figure 1C-D**). While no changes in cell morphology were observed at early timepoints (**Figure 1E-G**), at day 9 and 12 post-transduction, we found that double overexpression of OLIG2 and SOX10 partially changed the morphology of transduced astrocytes from non-ramified to OPC-like morphology (unipolar, bipolar or ramified) (**Figure 1G,H and Figure 2A,B**). No changes were observed in the conditions with single transduction or non-transduced control cells (**Figure 1E,F and H)**. Importantly, we found double-transduced cells expressing the oligodendrocyte marker O4, suggesting that astrocytes may be transitioning toward an oligodendrocyte lineage *in vitro* (**Figure 2C,D**). Based on these findings, we selected the combined overexpression of OLIG2 and SOX10 as the most effective condition to promote the transdifferentiation of human astrocytes into oligodendrocytes.

**Figure 2.**
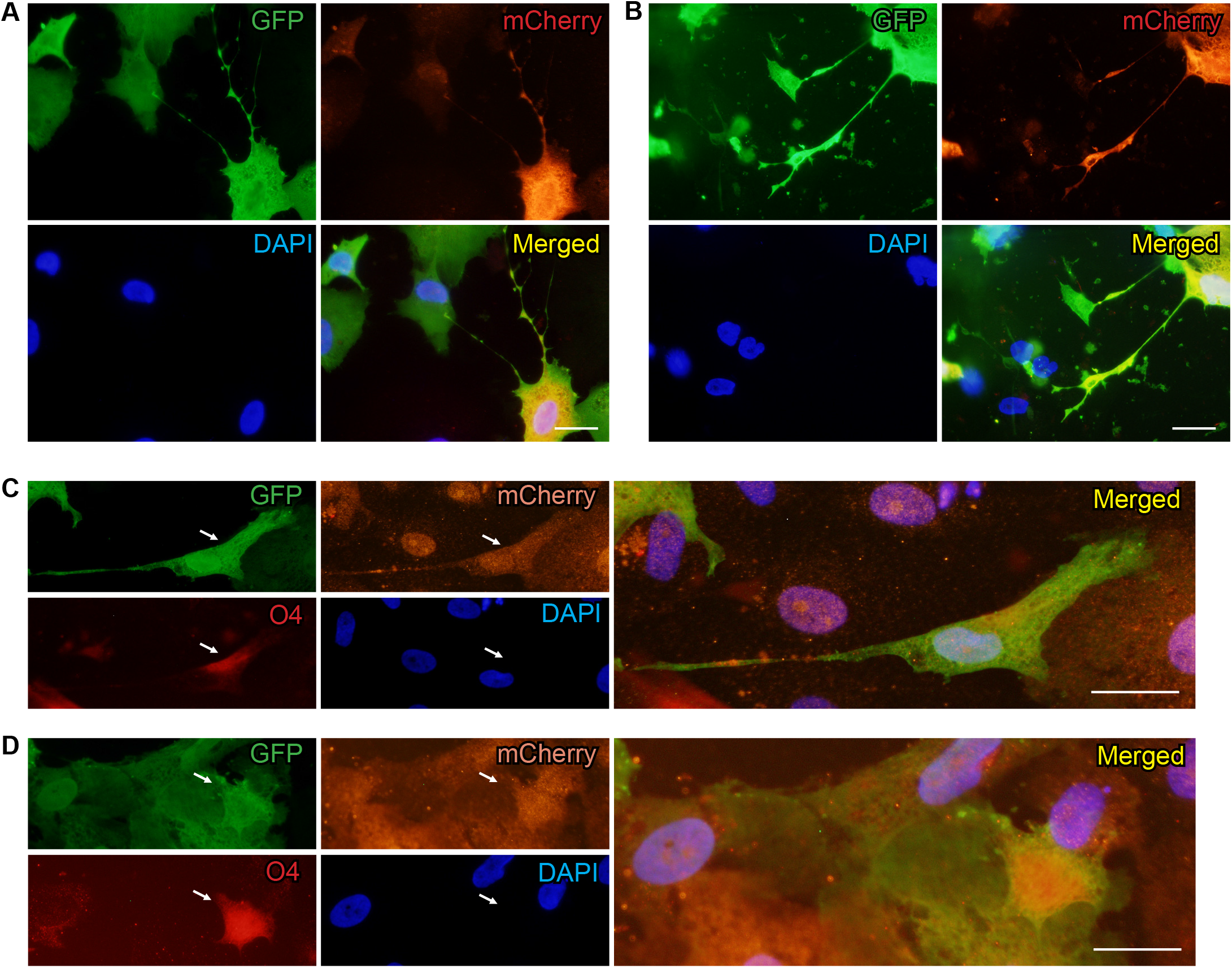
Morphological and molecular characterization of human converted oligodendrocytes 12 days post-transduction of human induced pluripotent stem (iPS) cell-derived astrocytes. (**A-B**) Representative immunofluorescence images of the cells generated 12 days post-transduction with GFAP-OLIG2-GFP and GFAP-SOX10-mCherry. Co-staining GFP and mCherry is shown. (**C-D**) Immunofluorescence images showing colocalization of GFP+mCherry+ converted cells with the oligodendrocyte marker O4. Arrows indicate colocalization. Scale bars, 10 μm.

### Human adult astrocytes successfully differentiate into mature oligodendrocytes in adult human cortical tissue *ex vivo*

Organotypic cultures of human adult cortical tissue provide a powerful platform for investigating cell reprogramming within a physiologically relevant environment that preserves human-specific cellular interactions and the three-dimensional architecture of the brain. We first assessed the preservation of the cortical tissue prior to reprogramming. In accordance with our previous findings (Palma-Tortosa et al. 2022; Gronning Hansen et al. 2020), we found expression of the neuronal markers NeuN and Map2, the astrocytic markers GFAP, Vimentin and S100β; and the oligodendrocyte markers OLIG2, SOX10, CNPase, and MBP (**Supplementary Figure 2A-C**).

To validate the cell-type specificity of lentiviral delivery in the human *ex vivo* system, tissue was transduced with either GFAP-OLIG2-GFP or GFAP-SOX10-mCherry (**Figure Supplementary 3A**). 3 days post-transduction, we observed that both GFP- and mCherry-transduced tissue displayed strong reporter expression localized to GFAP-positive glial cells, confirming selective targeting of astrocytes (**Figure Supplementary 3B**,**C**). Moreover, successful induction of OLIG2 and SOX10 expression was observed in GFP and mCherry-positive cells, respectively (**Figure Supplementary 3D**,**E**).

Based on our *in vitro* data, human cortical slices were transduced with both GFAP-OLIG2-GFP and GFAP-SOX10-mCherry lentiviral vectors to transdifferentiate astrocytes into oligodendrocytes. For reprogramming readout, the tissue was maintained for 12 days post-transduction. We observed broad expression of both GFP and mCherry reporters with a high percentage of colocalization (**Figure 3A**). Overexpression of both transcription factors, OLIG2 and SOX10 gave rise to cells with higher complexity compared to overexpression of only one transcription factor (**Figure 3B,C**). Importantly, we found double transduced cells expressing the mature oligodendrocyte marker CC1, arguing for the presence of mature reprogrammed-derived oligodendrocytes.

**Figure 3.**
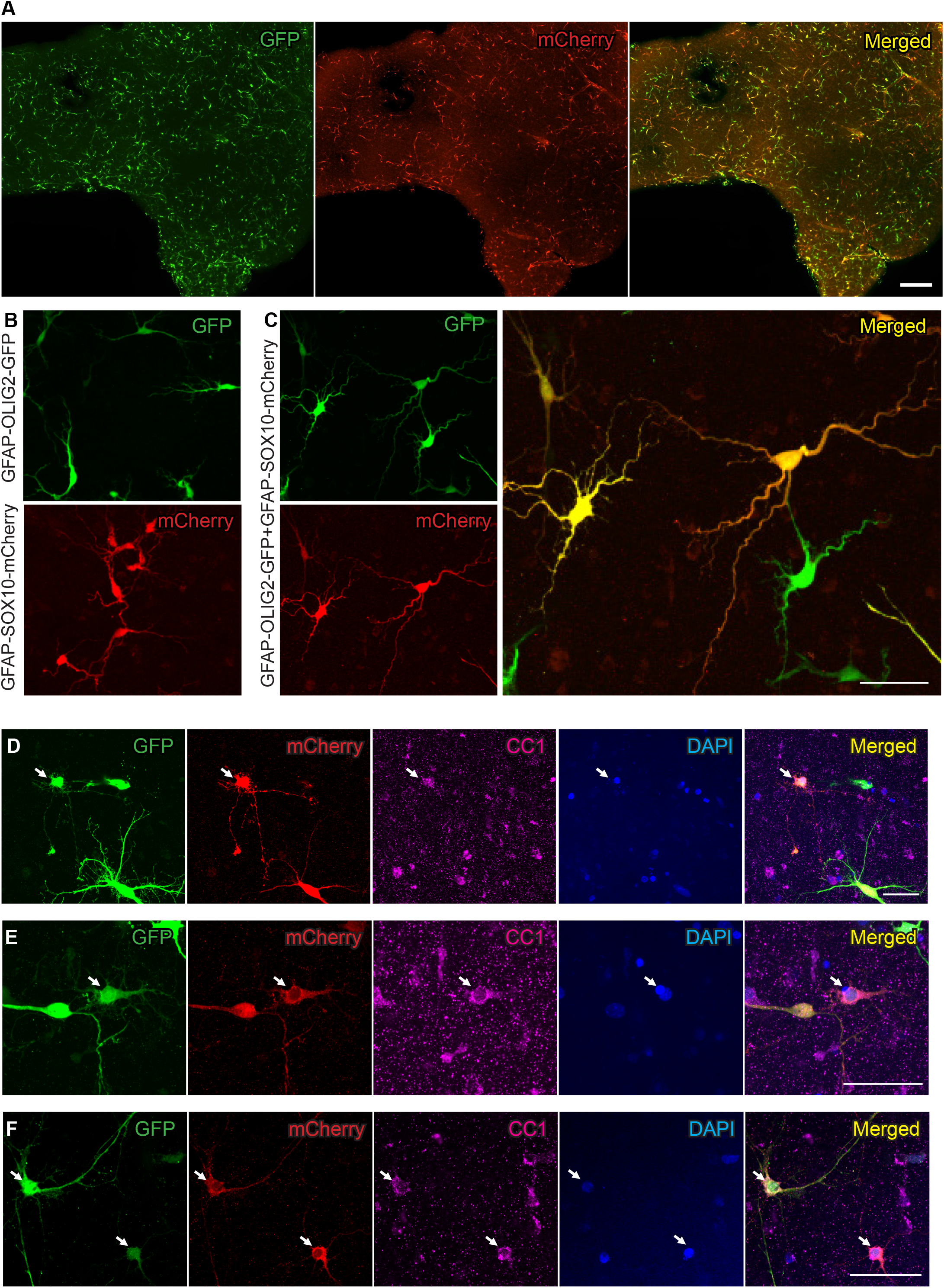
Overexpression of OLIG2 and SOX10 leads to transdifferentiation of astrocytes into mature oligodendrocytes in human adult cortical tissue. (**A**) Overview of reporters (GFP and mCherry) expression in organotypic cultures of human adult cortex 12 days after transduction with GFAP-OLIG2-GFP and GFAP-SOX10-mCherry lentiviruses. **(B-C)** Representative confocal images showing the morphological appearance of GFAP+ cells 12 days after overexpression of OLIG2, SOX10 (**B**) or a combination of both transcription factors (**C**). (**D-F**) Colocalizations of double transduced cells with the mature oligodendrocyte marker CC1. Arrows indicate co-expression of GFP, mCherry, CC1, and the nuclear marker DAPI. Scale bar in A, 500 μm. Scale bars in B-F, 50 μm.

To verify that the expression of CC1 is due to the transdifferentiation of astrocytes into mature oligodendrocytes and not because this marker is also expressed in the astrocytes existing in the human adult cortical tissue, we co-stained GFAP and CC1 in acute slices. No colocalizations were found (**Supplementary Figure 4A**). Moreover, no colocalizations with the neuronal marker NeuN were observed in GFP+/mCherry+ cells, indicating that no off-target neuronal populations were generated (**Supplementary Figure 4B**).

## DISCUSSION

Reprogramming of endogenous astrocytes into functional oligodendrocytes has been achieved in rodent models, as reported by several studies (Ghasemi-Kasman et al. 2018; Farhangi et al. 2019; Mokhtarzadeh Khanghahi et al. 2018)(Bajohr et al., 2025). However, whether this can be achieved in the human brain remains unknown. Here, we report that overexpression of the transcription factors SOX10 and OLIG2, which are involved in oligodendrocyte specification, is sufficient to generate mature oligodendrocytes from endogenous astrocytes, as shown by CC1 expression, in adult human organotypic cultures. To date, this is the first attempt to transdifferentiate astrocytes into oligodendrocytes in a human 3D environment.

*In vivo* animal studies involving transcription factor-mediated reprogramming were carried out by injecting viral vectors into mouse brains to induce astrocyte-to-oligodendrocyte transdifferentiation. Although the majority of the transduced cells were identified as astrocytes (75 to 95%), no specific promoter was used, and the origin of the remaining reprogrammed cells was not fully understood (Ghasemi-Kasman et al. 2018; Farhangi et al. 2019; Mokhtarzadeh Khanghahi et al. 2018). Here, we used the GFAP promoter to target reactive astrocytes, which has shown to be expressed in high levels in active MS lesions (Brosnan and Raine 2013; Rawji et al. 2020). This approach has been previously used for the conversion of mouse cortical astrocytes (Bajohr et al. 2025).

Here, we show that the overexpression of SOX10 and OLIG2 in GFAP-expressing endogenous astrocytes in human adult cortical tissue induce their transdifferentiation into mature oligodendrocytes in just 12 days, as observed by the expression of maturation marker CC1. Even if no conversion of human astrocytes into oligodendrocytes has been performed in human 3D systems, in vitro 2D conversion of the human astrocytoma line 1321N1 using the small molecules trichostatin A (TSA) or 5-azacytidine (Aza-C) gave rise to O4-positive cells at day 8 (Zare, Baharvand, and Javan 2019). Another combination of six small molecules (CHIR99021, Forskolin, RepSOX, LDN, VPA and Thiazovivin) was also found to be effective in producing OPCs from two human glioma lines (321N1 human astrocytoma and U87-MG human glioblastoma) (Sharifi-Kelishadi et al. 2024). In this study, the astrocytes gained OPC morphology 5 days after induction and expressed PDGFRα, O4 and MBP 15, 20 and 30 days respectively post-induction. Moreover, the transplantation of these chemically treated astrocytes into cuprizone-induced demyelinated mice brains eventually led to their conversion into mature oligodendrocytes 2 weeks after transplantation (Sharifi-Kelishadi et al. 2024; Zare, Baharvand, and Javan 2018). Even though these studies are of great interest, they were conducted using *in vivo* rodent models of demyelination, which overlook the critical human cell interactions.

To the best of our knowledge, this is the first study to evaluate direct lineage conversion of astrocytes into oligodendrocytes within a human tissue context that maintains the three-dimensional architecture of the patient’s brain. Future research will be necessary to determine the myelination capacity of the converted oligodendrocytes and to assess the potential of this strategy for structural and functional repair in the demyelinated human brain.

Clinical translation of reprogramming approaches remains an emerging and uncertain topic. Ongoing clinical trials in several neurodegenerative disorders (ie. Parkinson’s Disease, Alzheimer’s Disease or Huntington’s disease), where viral vectors are injected directly into the brain parenchyma, demonstrate that targeted gene delivery to the human CNS is technically feasible (Ye et al. 2024; Ling et al. 2023). This suggest that, with substantial further validation, the concept of reprogramming astrocytes into oligodendrocytes might one day be explored in a translational context for demyelinating disorders.

## METHODS

### Viral constructs

Lentiviral constructs were purchased from VectorBuilder Inc. To specifically target GFAP-expressing astrocytes, pLV-GFAP::hOLIG2-GFP and pLV-GFAP::hSOX10-mCherry were used.

### Generation of human iPS cell-derived astrocytes and *in vitro* reprogramming to oligodendrocytes

Human iPS cell-derived astrocytes were generated following the transcription factor–based protocol described by (Canals et al. 2018). Briefly, iPS cells were seeded on growth factor– reduced Matrigel–coated plates (Corning) and maintained in eTeSR medium (STEMCELL Technologies). The following day, cells were transduced with lentiviral vectors encoding reverse tetracycline-controlled transactivator (rtTA), SOX9, and NFIB. On day 0, transcription factor expression was induced by adding doxycycline (2.5 µg/mL; Sigma-Aldrich). On day 1, cultures were switched to expansion medium consisting of DMEM/F12 GlutaMAX (Thermo Fisher Scientific) supplemented with 10% FBS and 1% N2 (Thermo Fisher Scientific). From day 3 to day 5, cells were transitioned to FGF medium containing Neurobasal supplemented with 2% B27 (Thermo Fisher Scientific), 1% NEAA (Thermo Fisher Scientific), 1% GlutaMAX (Thermo Fisher Scientific), 1% FBS, 8 ng/mL FGF (Thermo Fisher Scientific), 5 ng/mL CNTF (Thermo Fisher Scientific), and 10 ng/ml BMP4 (Thermo Fisher Scientific). On day 7, cells were dissociated with Accutase, counted by trypan blue exclusion, and seeded onto the final experimental plates (Ibidi) at a density of 40.000 cells per well. On day 14 (refered as 0 days post-transduction or 0 DPT), astrocytes were transduced with lentiviral vectors encoding OLIG2 and SOX10 in glial induction medium, composed of DMEM/F12 supplemented with 1% B27 without vitamin A, 0.5% N2 supplement, and 1% GlutaMAX. Starting two days post-transduction, cultures were supplemented with T3 (60 ng/mL), PDGF (10 ng/mL), and SAG (1 µM). Media was refreshed every 2–3 days, and cells were maintained up to 12 days post-transduction. Images of live cells were captured every third day using fluorescent microscope (IX51, Olympus, Germany).

### Immunocytochemistry

*In vitro* astrocytes were fixed 12 days post-transduction in 4% PFA for 10 min. This was followed by washing three times with KPBS for 10 min each. They were incubated with blocking solution (1% Triton X-100 in KPBS and 5% NDS) for 30 min. Primary antibodies (**Table 1**) diluted in blocking solution were added to the astrocytes and incubated for 3 h. Cells were then washed three times for 10 min each with KPBS. Secondary antibodies were added for 45 min, followed by three rinses with KPBS. Then DAPI was added for 10 mins, and then washed with KPBS. All staining steps were carried out at room temperature. For O4 staining, the blocking solution contained KPBS and 5% NDS, non-detergent added. Images were captured using either an epifluorescent microscope (IX73, Olympus, Germany) or a confocal microscope (LSM 780, Zeiss, Germany) or Olympus IX73 microscope.

**Table 1.** List of primary and secondary antibodies (in alphabetical order) used for immunohistochemistry and immunocytochemistry. AR: Antigen retrieval

| Antibody | Company (Catalogue No.) | Dilution | Notes |
| --- | --- | --- | --- |
| CC1 (mouse IgG) | Abcam (ab16794) | 1:500 | AR |
| CNPase (rabbit IgG) | Abcam (ab227218) | 1:500 | AR |
| Donkey anti-chicken IgG | ThermoFisher/Alexa fluor 488 | 1:500 |  |
| Donkey anti-chicken IgG | ThermoFisher/Alexa fluor 647 | 1:500 |  |
| Donkey anti-mouse IgG | ThermoFisher/Alexa fluor 488 | 1:500 |  |
| Donkey anti-mouse IgG | ThermoFisher/Alexa fluor 568 | 1:500 |  |
| Donkey anti-mouse IgG | ThermoFisher/Alexa fluor 647 | 1:500 |  |
| Donkey anti-rabbit IgG | ThermoFisher/Alexa fluor 488 | 1:500 |  |
| Donkey anti-rabbit IgG | ThermoFisher/Alexa fluor 568 | 1:500 |  |
| Donkey anti-rabbit IgG | ThermoFisher/Alexa fluor 647 | 1:500 |  |
| Goat anti-mouse IgM | Invitrogen (A21238) | 1:500 |  |
| GFAP (chicken IgG) | Abcam (ab13970) | 1:1000 |  |
| GFP (chicken IgG) | Millipore (AB18901) | 1:1000 |  |
| MAP2 (chicken IgG) | Abcam (ab5392) | 1:1000 |  |
| MBP (mouse IgG) | Biolegend (808401) | 1:1000 | AR |
| mCherry (mouse IgG) | Abcam (ab125096) | 1:500 |  |
| NeuN (rabbit IgG) | Abcam (ab104225) | 1:500 |  |
| O4 (mouse IgM) | R&D Systems (MAB1326) | 1:200 |  |
| OLIG2 (rabbit IgG) | Abcam (ab109186) | 1:500 | AR |
| RFP (chicken IgG) | Rockland (600901379) | 1:500 |  |
| RFP (mouse IgG) | Abcam (ab65856) | 1:500 |  |
| RFP (rabbit IgG) | Abcam (ab62341) | 1:500 |  |
| S100B (rabbit IgG) | DAKO (GA504) | 1:500 | AR |
| SOX10 (rabbit IgG) | Abcam (ab180862) | 1:500 | AR |
| STEM123 (mouse IgG) | Takara (AB-123) | 1:1000 |  |
| Vimentin (mouse IgG) | Abcam (ab24525) | 1: 500 | AR |

### Collection and processing of human adult cortical tissue

All the experiments concerning the use of human tissue were carried out under the guidelines approved by the Regional Ethical Committee, Lund, Sweden (ethical permit number 2021-07006-01). Human adult cortical tissue was obtained with informed consent from patients undergoing elective surgery for temporal lobe epilepsy. Prior to the day of the surgery, cutting solution, rinsing solution, and human tissue media were prepared as described previously (Palma-Tortosa et al., 2022). The samples were brought to the lab immediately after re-sectioning from the patient in the operation room. The tissue was cut coronally with a thickness of 300 µm in a vibratome (Leica, VT200S).

### Maintenance of organotypic cultures of human adult cortical tissue

For tissue culture, the samples were transferred to culture inserts (Millipore, PICMORG50) and cultured in previously equilibrated human tissue media, containing Brain Phys medium [without phenol red] supplemented with B27 [1:50], glutamax [1:200], antibiotic-antimycotic [1:50] and gentamycin [50 mg/mL]. Tissue slices were cultured in an incubator (37 °C, 5% CO_2_ and 90-95% humidity). The media was changed every other day supplemented with BDNF (50 ng/ml, Peprotech) and NT3 (50 ng/ml, Peprotech). For further details see (Palma-Tortosa et al. 2022).

### *Ex vivo* reprogramming

Viruses were diluted in human tissue media at a ratio of 1:100, and the solution was added on the top of semi-dry tissues. 24 h post-transduction, 300 µL of media was added on top of the insert to prevent the tissue from drying out. On the following day, the media was removed and fresh human tissue media was added. Media was changed every other day.

### Immunohistochemistry

Human adult cortical tissue slices were fixed two weeks post-transduction in 4% paraformaldehyde (PFA) overnight at 4 °C on a shaker, followed by three 15-minute rinses with potassium-containing phosphate-buffered saline (KPBS). Next, slices were incubated overnight at 4 °C in permeabilization solution (0.02% Bovine Serum Albumin (BSA) and 1% Triton X-100 in KPBS) and the following day with blocking solution (KPBS with 0.2% Triton X-100, 1% BSA, sodium azide [1:10,000], and 10% normal donkey serum (NDS) (Merck Millipore)) for at least 4 h at 4 °C. Primary antibodies (**Table 1**) were diluted in blocking solution and incubated for 48 h at 4°C. Fluorophore-conjugated donkey secondary antibodies (**Table 1**), also diluted in blocking solution, at a ratio of 1:100 and were applied for 24 h at 4 °C. Following this, samples were washed and stained with 4′,6-diamidino-2-phenylindole (DAPI) for 2 h at room temperature. Finally slices were rinsed with KPBS, mounted on glass slides with Dabco (Merck) mounting media and coverslipped with 1.5 mm thickness. Stained slides were scanned on Virtual Slide Scanning System (VS-120-S6-W, Olympus, Germany) to get an overview of the marker expression. High magnification images were captured using a confocal microscope (LSM 780, Zeiss, Germany).

Some epitopes (see **Table 1**) required antigen retrieval. Therefore, the above staining protocol was modified to perform DAPI staining prior to permeabilization step. Next, antigen retrieval was performed using sodium citrate pH 6.0, Tween 0.05%, for 2 h at 65 °C. For samples undergoing antigen retrieval, 1% Triton X-100 was used in blocking solution.

## AUTHOR CONTRIBUTIONS

S.P.-T., conceived the project. A.P., L.R. R.M-C., K.E.O., W.G.E., and S.P.-T., conducted the experiments and analyzed the data. J.B., H.A., provided human material. A.P., and S.P.-T., wrote the manuscript. A.P., and S.P.-T., were involved in collecting and/or assembly of data, data analysis, and interpretation. All authors reviewed and edited the manuscript.

## ACKNOWLEDGMENTS

This work is supported by grants from the Crafoordska Stiftelsen (20250660), Åke Wibergs Foundation (M24-0186 and M25-0303), Neurofonden, Rut och Erik Hardebo Foundation, and the Thorsten and Elsa Segerfalk Foundation to S.P-T. Authors acknowledge the Lund Stem Cell Center’s Cell and Gene Therapy Core for the technical support.

## CONFLICT OF INTERESTS

The authors declare no competing interests.

## Notes

### Competing Interest Statement

The authors have declared no competing interest.

