## Supplementary data for "*Ex vivo* astrocyte-to-oligodendrocyte conversion in human adult cortical tissue using transcription factor overexpression"

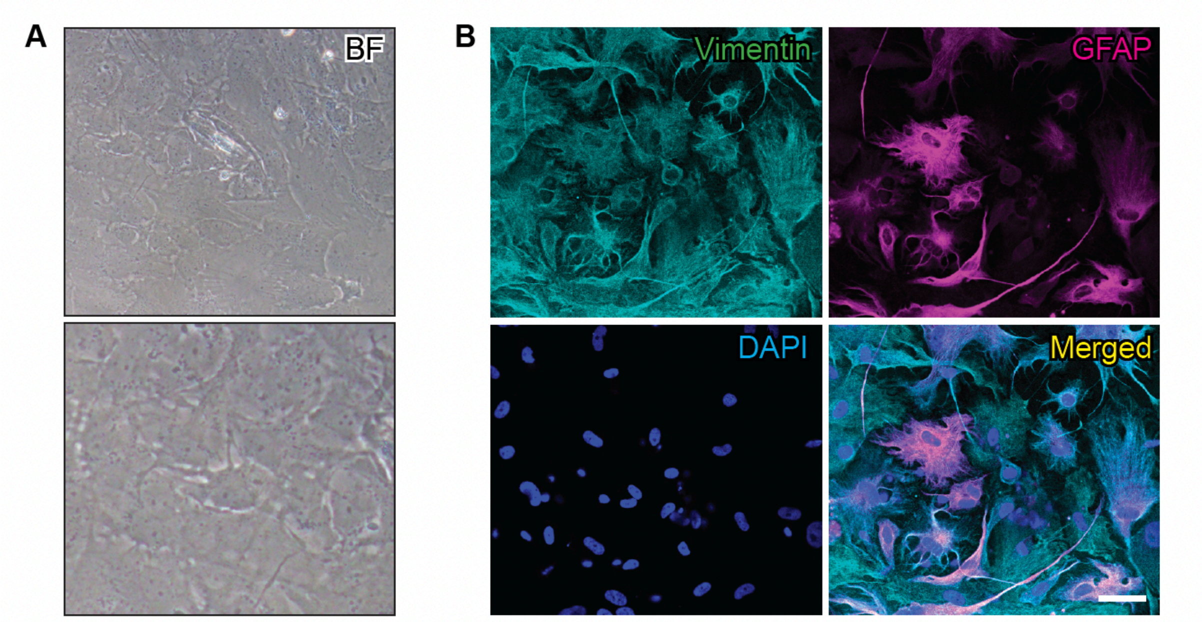


Supplementary Figure 1. **Morphology and molecular expression of human induced pluripotent stem (iPS) cell-derived astrocytes.** Representative brightfield images (**A**) and immunocytochemistry (**B**) showing expression of the astrocytic markers vimentin and glial fibrillary acidic protein (GFAP). Scale bar, 50 µm.

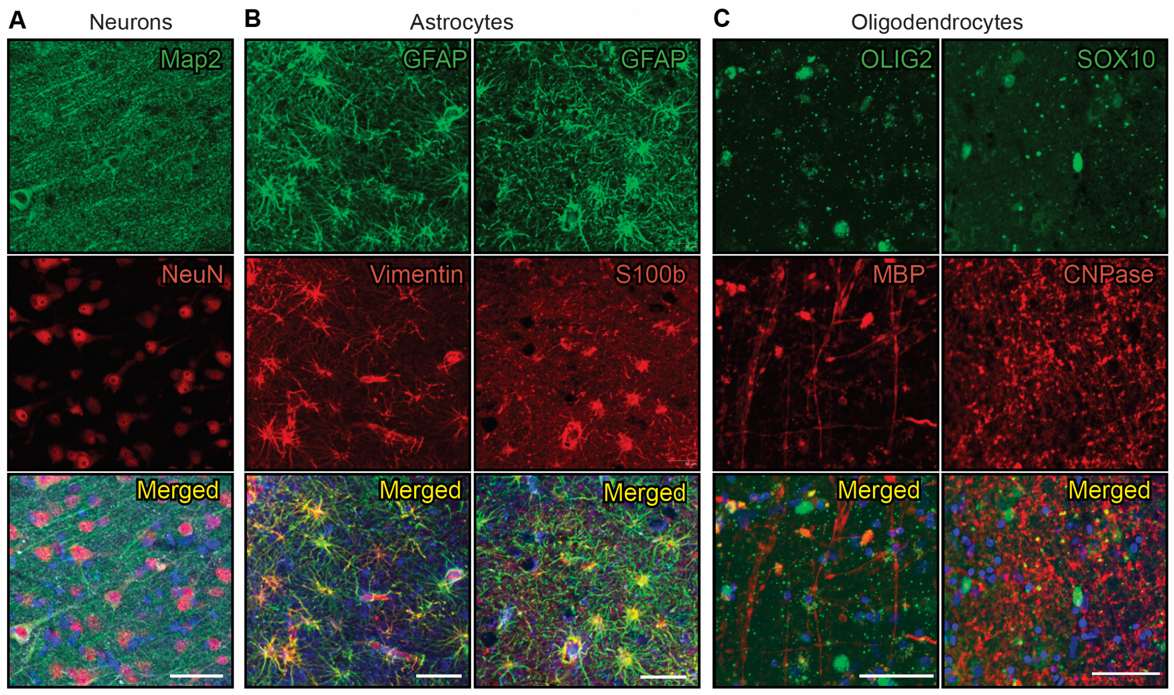


Supplementary Figure 2. **Characterization of the neural cell populations in acute human adult cortical tissue**. Representative confocal images showing the expression of the neuronal markers Map2 and NeuN (**A**), the astrocyte markers GFAP, Vimentin and S100β (**B**) and the oligodendrocyte markers OLIG2, SOX10, MBP, and CNPase (**C**). Nuclear staining (DAPI, blue) is included in the merged panel. Scale bars, 50 μm.

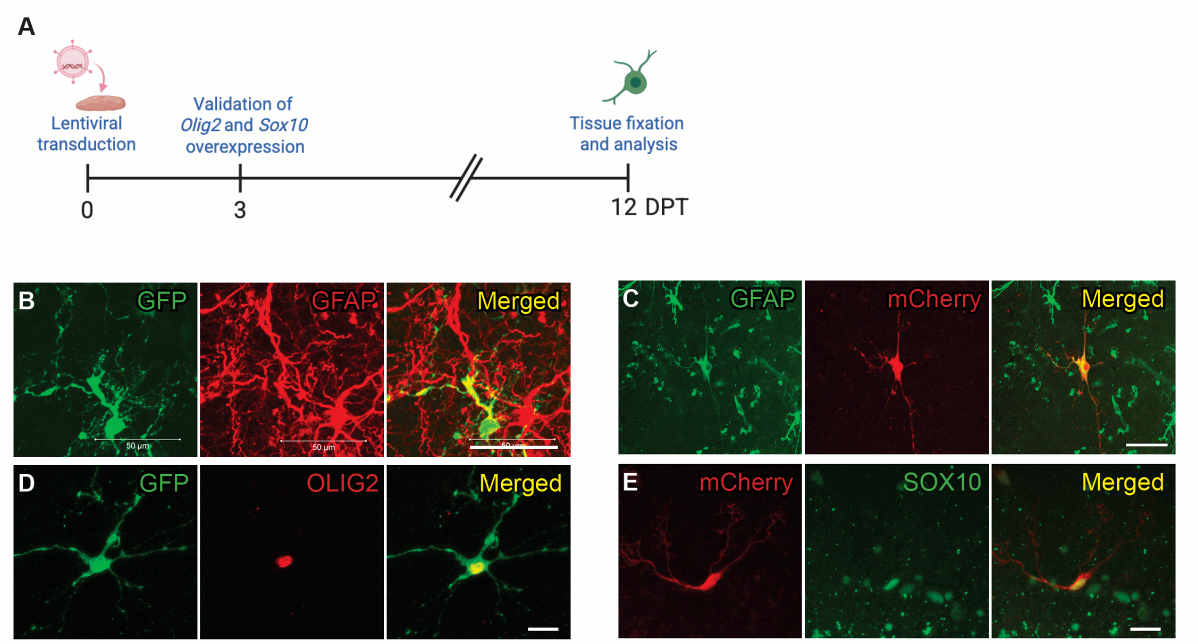


**Supplementary Figure 3. Validation of lentiviral-mediated overexpression of OLIG2 and SOX10 in human adult cortical tissue 3 days post-transduction. (A)** Experimental workflow for the transdifferentiation of GFAP-expressing astrocytes into oligodendrocytes in organotypic cultures of human adult cortex. (**B**-**E**) Representative confocal images showing colocalization of the astrocyte marker GFAP with the reporters GFP (**B**) and mCherry (**C**). Colocalizations of GFP with OLIG2 (**D**) and mCherry with SOX10 (**E**). Scale bars in B and D, 50 µm. Scale bars in C and E, 20 µm.

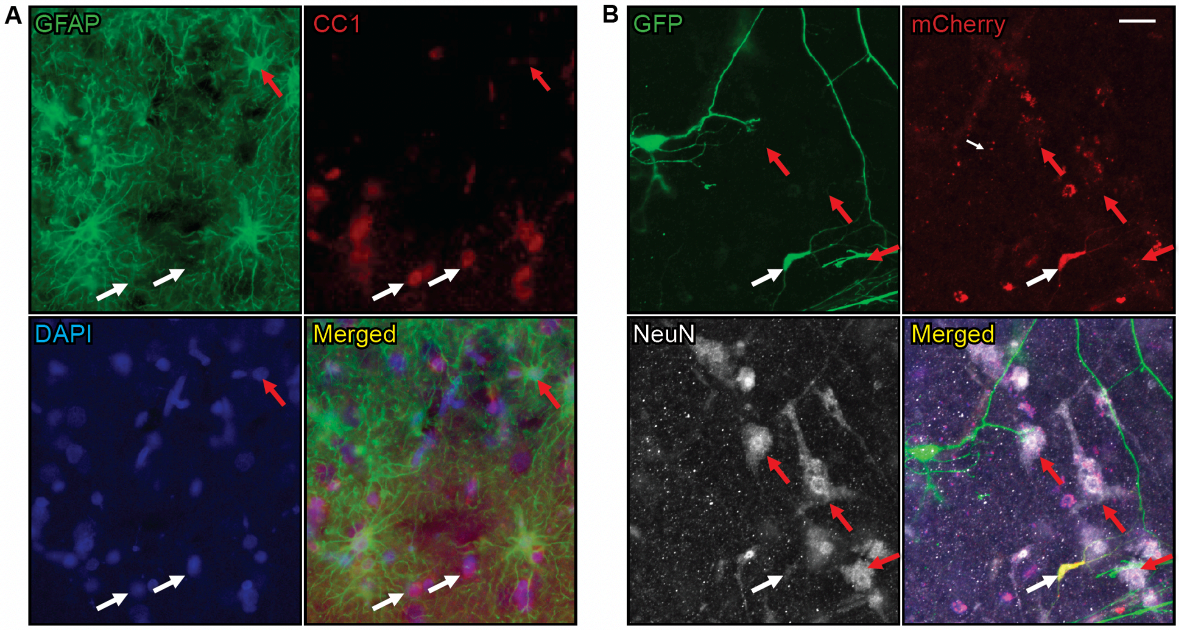


Supplementary Figure 4. **Overexpression of OLIG2 and SOX10 in GFAP-expressing human astrocytes does not induce neuronal differentiation.** (**A**) Representative images showing lack of colocalization of the astrocyte marker GFAP and the mature oligodendrocyte marker CC1. (**B**) Representative images showing lack of colocalization of the double transduced cells (GFP+mCherry+) and the neuronal marker NeuN. White arrows indicate CC1+ cells (in **A**) and a GFP+mCherry+ cell (in **B**). Red arrows indicate a GFAP+ cell (in **A**) and NeuN+ cells (in **B**). Scale bars, 20 µm.
